# The genome wide analysis deciphered virulence factors and secondary metabolites of *Chryseobacterium* and description of *Chryseobacterium indicum* sp. nov

**DOI:** 10.1101/2022.03.19.484977

**Authors:** Manish Prakash Victor, Lipika Das, Subrata K. Das

## Abstract

This study describes a novel bacterium, *Chryseobacterium* strain PS-8, isolated from the skin of freshwater pufferfish (*Tetraodon cutcutia*). Cells are Gram-negative. The genome size is 4.72 Mb, and the G+C content is 36.4%. The *in silico* DDH homology, ANI, and AAI were below the cutoff value, 70% and 95% to 96%, respectively, as required for novel species delineation. The phylogenomic analysis using core and non-recombinant core genes, the strain PS-8 showed clustering with *Chryseobacterium gambrini* DSM 18014^T^. Generally, *Chryseobacterium* species are considered opportunistic pathogens. Prediction of the virulence genes revealed genes for adherence, biofilm and stability *(glf, lpxD, clpE, cps0, clpE, IlpA*), proliferation (*tufA, cap8E, galE, kfic, bioB, clpP, kdtB, csgD, carB*), resistance to immune response (*htpB, katA, wbtl, sodB, kpsF*), and host-defense evasion system (*kpst, clpE, kfic, ybtQ, cap8G, clpP*). The cladogram based upon the virulence genes showed a similar phylogeny amongst the *Chryseobacterium* species. Additionally, secondary metabolites producing gene clusters were identified, including microviridin, resorcinol and polyene, terpene, etc. Our study showed that strain PS-8 constitute a novel species for which *Chryseobacterium indicum* sp. nov. is proposed. The type strain is PS-8T (= TBRC 15233^T^ = NBRC 115235^T^).

## INTRODUCTION

The genus *Chryseobacterium* was described by Vandamme *et al.* (1994). Many species of the genus *Chryseobacterium* are described from various environments, including soil, phyllosphere, fish, industrial plants and clinical samples (Behrendt *et al.* 2007; Herzog *et al.* 2008; Yassin *et al.* 2010; Li and Zhu H 2012; Zamora *et al.* 2012). Despite being a biocontrol agent for plant pathogens (Kim *et al.* 2012), it can cause severe infections in immune-compromised humans (new born and ≥65 years). Several *Chryseobacterium* isolates were reported from hospitalised patients respiratory tract or blood. Also, some species of *Chryseobacterium* have been described as causative agents for diseases like emphyema, pneumonia and meningitis (Bloch *et al.* 1997) and act as sporadic but severe opportunistic nosocomial pathogens (Hsueh *et al.* 1997; Chiu *et al.* 2000). The significant species isolated in infected patients are *C. indologenes* (Izaguirre-Anariba and Sivapalan 2020) and *C. gleum* (Tsouvalas *et al.* 2020). In addition, certain strains show matrix digesting properties such as collagenous matrices, feathers and exoskeletons (Page *et al.* 2019). *Chryseobacterium* appeared as yellowish-orange colonies because of a pigment flexirubin. This pigment has biotechnological application. It can be used as additives or supplements in cosmetics, pharmaceuticals, livestock feed and food industries (Kirti *et al.* 2014; Venil *et al.* 2014). The genus *Chryseobacterium* is Gram-negative, non-spore-forming, aerobic nonfermenting rods-shaped bacteria. It is non-motile. Growth temperature ranges from 5° - 30° C, but strains isolated from human specimens grow at 37 °C. G + C mol% of DNA range from 29 - 39% (Bernardet *et al.* 2010). The Major fatty acids are iso-C15:0, iso-C17:0 3OH, iso-C15:0 3OH and iso-C15:0 2-OH and polar lipids include phosphatidylethanolamine, amino lipids and phospholipids (Kämpfer *et al.* 2009).

When writing, the genus *Chryseobacterium* included 127 validly published named species as of 5 November 2021 (https://lpsn.dsmz.de/genus/chryseobacterium). In this regard, we compared the genome of the strain PS-8 with 73 genome sequences of the type strains of the genus *Chryseobacterium* available in the NCBI database. The phylogeny based on the genome-wide core genes and evaluation of ANI and *in silico* DDH has been used to ascertain the novelty of the strain PS-8 (Mateo-Estrada *et al.* 2019). This study described the genomic comparison of strain PS-8 with validly published named species of the genus *Chryseobacterium* to delineate the taxonomic assignments. Additionally, we predicted the genes encoding virulence properties and mapped the secondary metabolites in this group of bacteria.

## RESULTS AND DISCUSSION

### Phenotypic characterization

The **s**train PS-8 appeared as yellowish-orange, smooth colonies, up to 1.0 mm in diameter. The strain was Gram-reaction-negative, strictly aerobic, with rod-shaped cells of approximately 0.5–0.75 μm wide and 0.8 - 2.0 μm long in size (**Figure S1**). Growth temperature in nutrient broth ranges from 10-40 °C with optimal growth at 28–37 °C. The optimal pH for growth is between 4.5–11, and the optimum growth is at pH 5.5–7.5. Strain PS-8 could grow in the presence of 0–6 % NaCl (w/v), and the optimal growth occurred at 0.5% NaCl. The strain PS-8 was negative for haemolytic activity. The different growth responses of the strain PS-8 are shown in (**Table S1**). The *Chryseobacterium* type species reported worldwide as of 5 November 2021 has mentioned in **Fig. 1.**

**Fig. 1.**
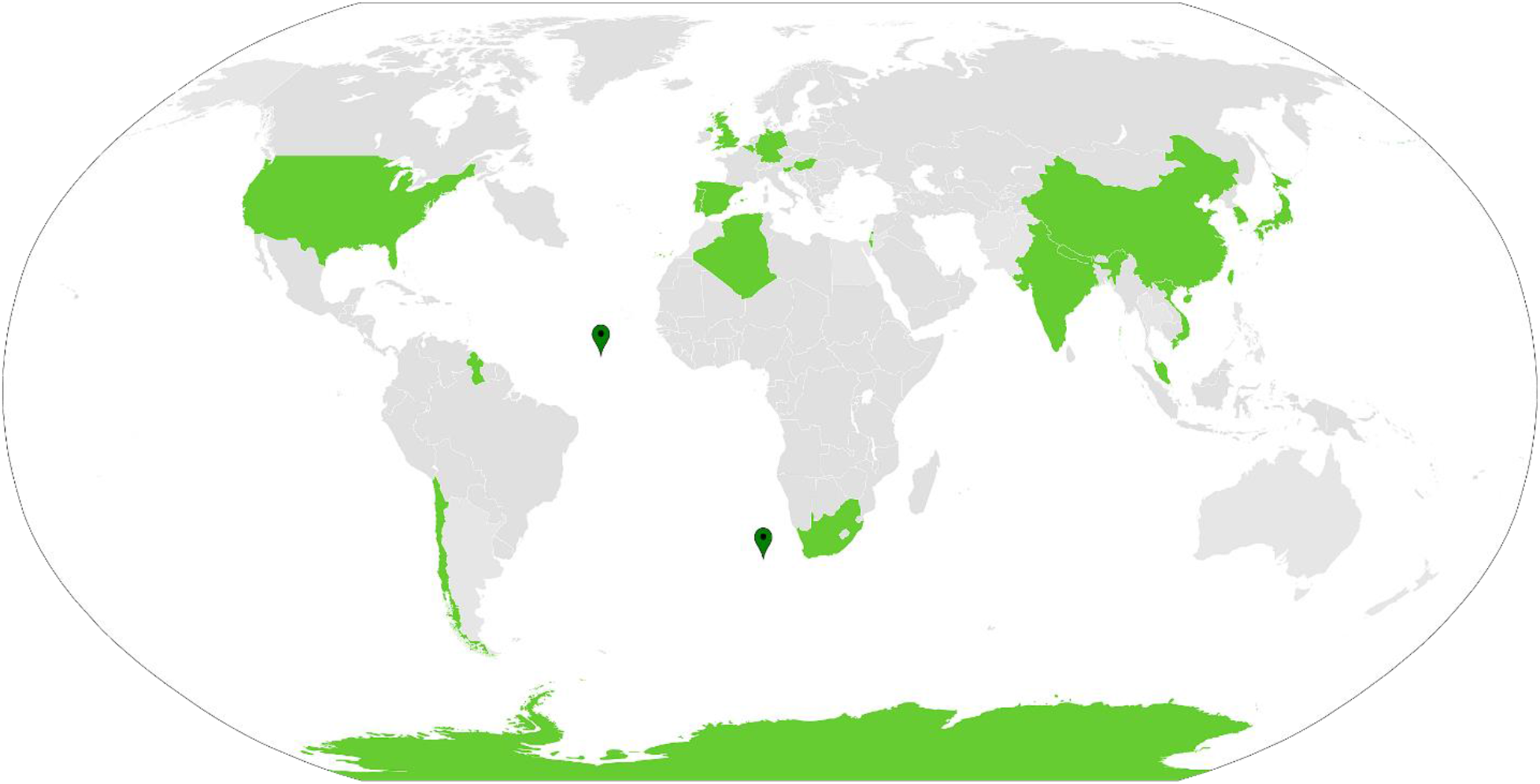
*Chryseobacterium* type species reported as on 5th November, 2021 world-wide. Korea, (35); China, (19); Germany, (11); Spain, (9); USA, (8); Taiwan, (6); South Africa, (5); Israel, (4); India, (4); Antarctica, (4); Japan, (3); Artic, (2); Belgium, (2); Chile, (2); Portugal, (2); Nepal, (2); Algeria, (1); Guyana, (1); Scotland, (1); Malaysia, (1); Slovenia, (1); Hungary, (1); Vietnam, (1); North Atlantic ocean, (1); South Africa Atlantic ocean, (1).

### Chemotaxonomic characterization

The Polar lipids included phosphatidylethanolamine (PE), four amino lipids (AL1-4), and nine unknown lipids (L1-9) (**Figure S2**). The lipid profile of strain PS-8 was closely related to the reference strains of the genus *Chryseobacterium* (Montero-Calasanza *et al.* 2014). The strain PS-8 contained 15:0 iso (38.04%), 17:0 iso 3OH (16.51%), 15:0 iso 3OH (3.15%), and 11:0 anteiso (2.38%) as the major fatty acids (**Table S2**). The composition of fatty acids is consistent with the type species, justifying its placement in the genus *Chryseobacterium* (Kämpfer *et al.* 2009). The cell wall sugars contained glucose, galactose and ribose. The meso-DAP was found the type of diaminopimelic acid. The overall chemotaxonomic characteristics of strain PS-8 support its placement to the genus *Chryseobacterium.*

### Genome features

The genome of the strain PS-8 was 4.72 Mb, and the G+C content was 36.4%. The de novo assembly produced five scaffolds covering 4,743,343 bp (85.983%). The genome had 4397 genes, including 4251 protein-coding genes (CDS). The genome contains 90tRNA, 5 5SrRNA, 5 16S rRNA, and 5 23S rRNA. Gene ontology study exhibited (45.02%) genes were associated with biological processes (45.02%), followed by (37.99%) with molecular function and (16.92%) cellular components (**Figure S3**). The GenBank/EMBL/DDBJ accession number for the genome and 16S rRNA gene sequences of *Chryseobacterium indicum* strain PS-8 are <u>JACSGT000000000</u> and <u>MZ305332</u>, respectively.

### Overall genome-related indices and phylogenetic analysis

The whole-genome sequence information is essential for comparative phylogenetic and taxonomic studies of microorganisms. In this regard, we compared the genomic relatedness of the strain PS-8 with the genomes of the *Chryseobacterium* type strains (NCBI, November 5, 2021). The ANI and AAI values of the strain PS-8 with the *Chryseobacterium* type strains were below (95-96%), suggesting a novel species (Konstantinidis and Tiedje 2005a; Konstantinidis and Tiedje 2005b). Further, isDDH similarity values were less than 70 % to define bacterial species (Meier-Kolthoff *et al.* 2013). These results indicate the strain PS-8 is a novel species in the genus *Chryseobacterium* (**Table 1; Table S3**). Further, we analyzed the phylogenetic relationship between the strain PS-8 and the reference strains based on core and non-recombinant core genes. The maximum-likelihood (ML) phylogenetic tree showed the strain PS-8 clustered with *Chryseobacterium gambrini* DSM 18014^T^, and it formed a distinct clade (**Fig. 2**) which is similar to the phylogenetic tree based on the non-recombinant genes (**Figure S4**), indicating the robustness of the tree topology.

**Table 1.**
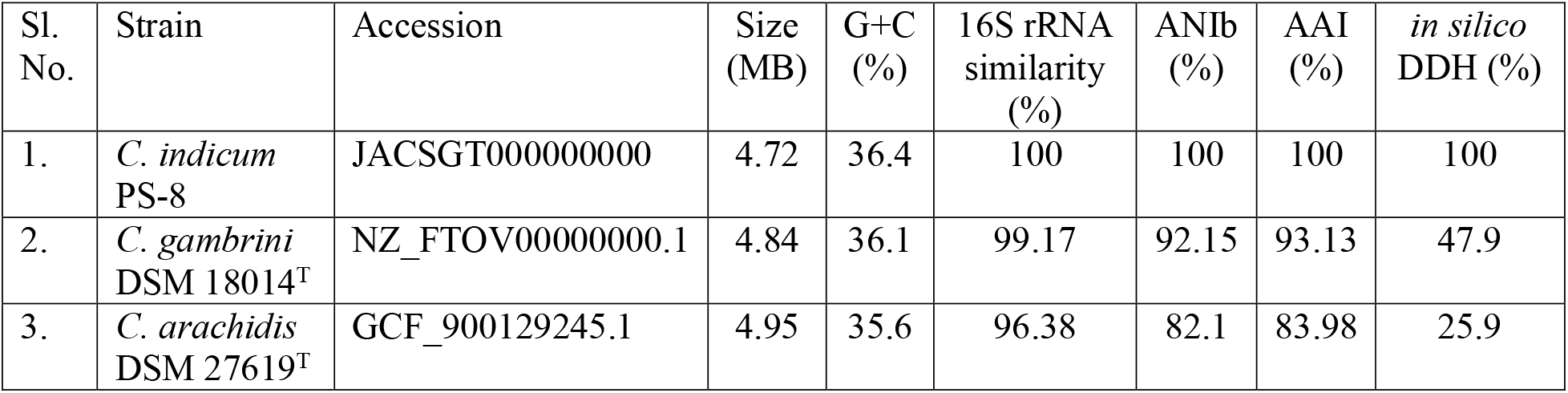
Comparison of the genomic characteristics of *Chryseobacterium* strain PS-8 with closely related species of *Chryseobacterium*.

**Fig. 2.**
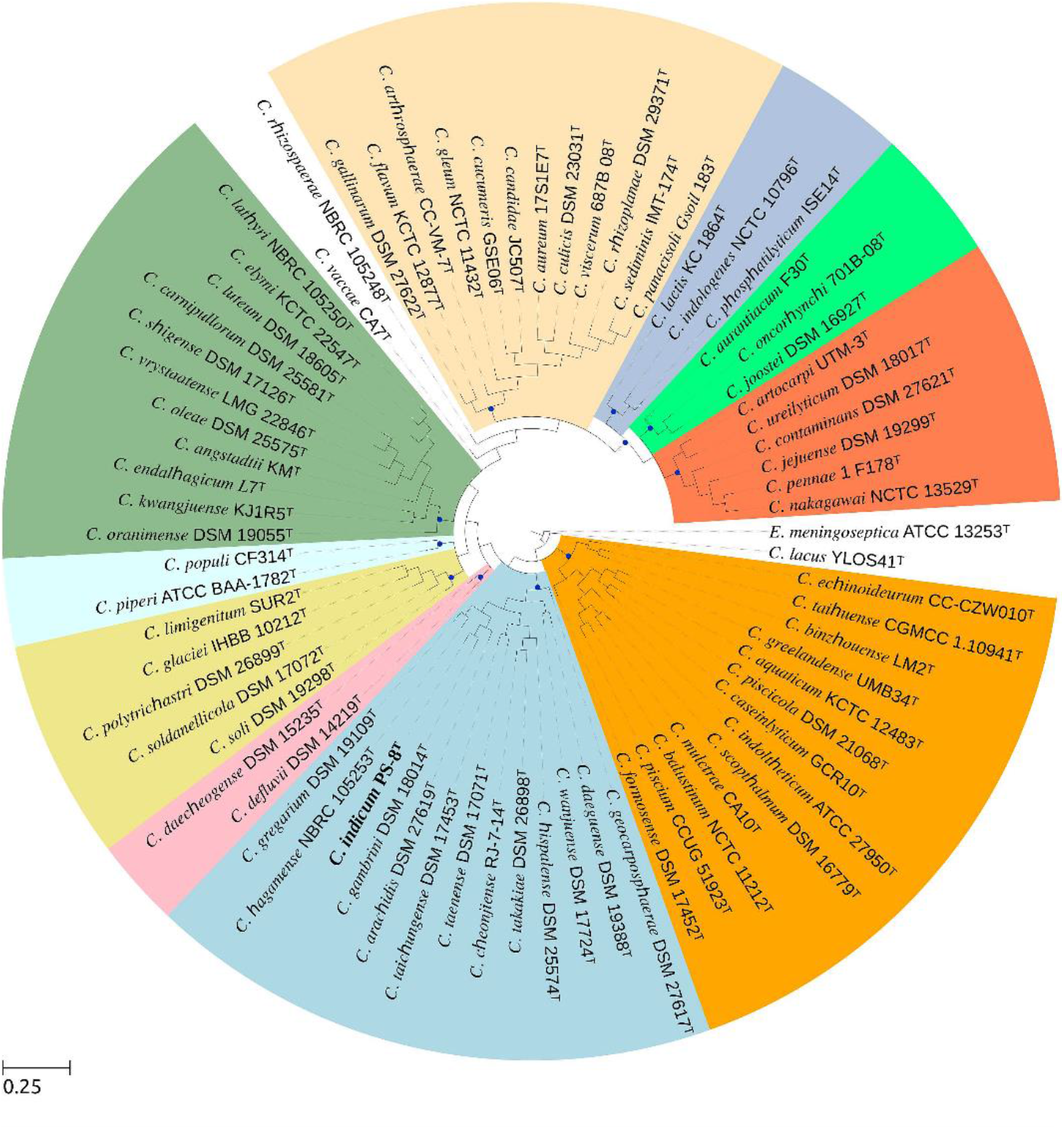
Maximum-likelihood phylogenetic tree based on core gene sequences of *Chryseobacterium* strain PS-8 with available genome sequences of 73 type species. Bootstrap values > 62% for all the branches based on 1000 resampling. Bar, 0.25 substitutions per nucleotide position.

### Analysis of virulence genes

In total, 41 virulence-related genes were predicted in the genome of the strain PS-8 using the CG view. The rings 1 and 2 indicated with green arrows in both forward and reverse direction demonstrates the protein-coding genes (CDS). Additionally, the zoomed portion of the image suggests the presence of genes encoding virulence factors (**Fig. 3**). The virulence genes identified showed closeness to genes that reveals functions like (i) adherence, biofilm and stability (*glf, msrA/B_pilB, acpxL, lpxD, clpE, cps0, clpE, IlpA*); (ii) proliferation (*tufA, cap8E, icl, manC, fleQ/flrc, galE, kfic, bioB, clpP, kdtB, csgD, carB*); (iii) resistance to immune response (*htpB, katA, wbtl, cj1136, hasB, kdsA, lirB, gnd, clbC, cylG, sodB, kpsF*) and (iv) host-defense evasion (*kpst, clpE, kfic, ybtQ, cap8G, clpP, FTT-0798*) (**Figure S5**). The functions of the virulence genes and the sessile nature of the bacterium could play an essential role in its survival and spread. Since the information on the pathogenesis of *Chryseobacterium* spp is limited, this study hypothesized that attachment of the bacterium to the host cells/organs could be accomplished through the gene described above set (i). After the invasion into the host, the gene set (ii) and (iii) might come into the picture for the sustained growth and expansion of the bacterial colonies. Eventually, the gene set (iv) become active in establishing the bacteria in the host. Based on the (40%) identity cut-off, 11 virulence genes (*clpP, hasB, acpxL, htpB, kdsA, tufA, katB, wbtl, galE, kpsE, cpsA)* from the strain PS-8 were selected for comparison with the 73 type species of *Chryseobacterium.* The cladogram and the heatmap based upon the virulence genes showed the closeness among the *Chryseobacterium* species (**Fig. 4**). Thus, the virulence genes predicted in this study provide a preliminary insight into the pathogenicity of *Chryseobacterium* and warrant further studies.

**Fig. 3.**
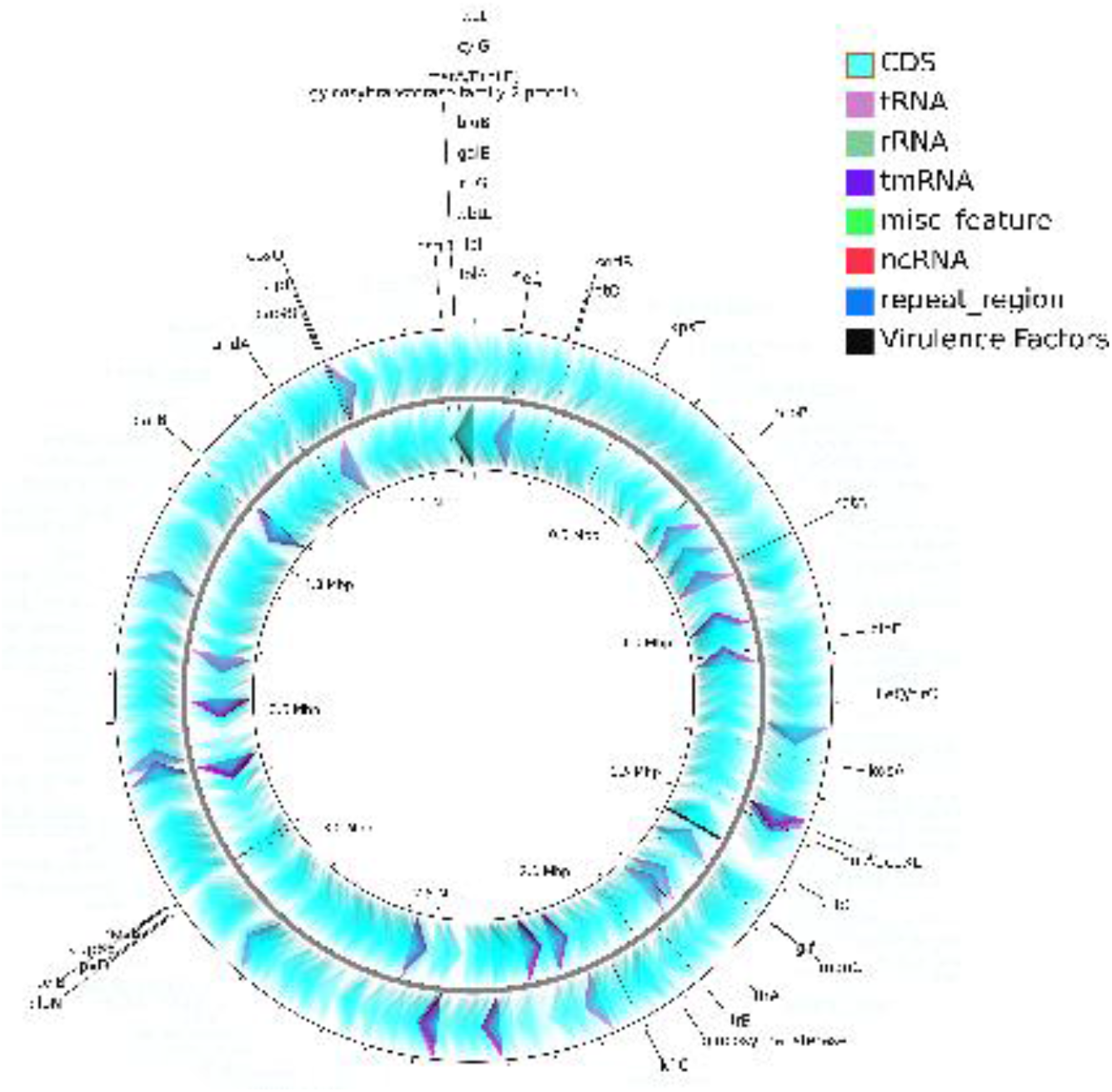
The CG view server BETA created a circular map of the *Chryseobacterium* strain PS-8 genome.

**Fig. 4.**
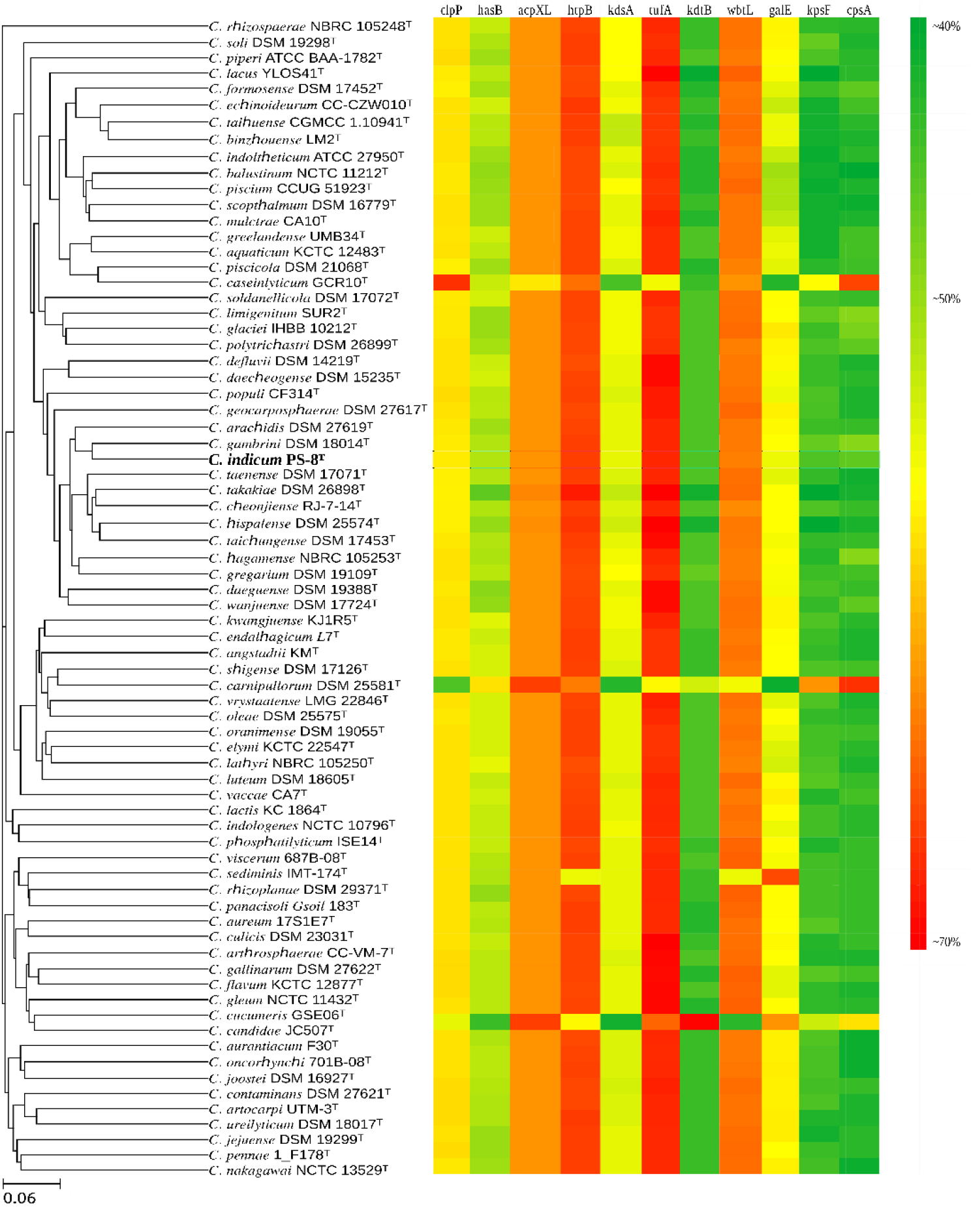
Cladogram-heatmap based upon protein sequences of virulence genes in *Chryseobacterium*. Conservation in nucleotide sequences of orthologous virulence genes in *Chryseobacterium* strain PS-8 with reference strains. Bar represents percent nucleotide sequence identity.

### Prediction of genes encoding secondary metabolites

We investigated the genes encoding for the putative biosynthetic gene clusters (BGCs) in the genus *Chryseobacterium*. In this regard, the *Chryseobacterium* type strains with available whole-genome sequences were grouped based on the niche categories, and we predicted the genes encoding secondary metabolites produced by them. The result shows the metabolites like microviridin, aryl polyenes-resorcinol and terpenoids gene clusters present in the strain PS-8. The genomic context of these genes identified in the strain PS-8 compared with that in the related *Chryseobacterium* species obtained through antiSMASH. We Found the maximum similarity with aryl polyenes-resorcinol (80%) followed by terpene (28%) and microviridin (**Fig. 5**). In addition, metabolites like siderophore, ladderanes, proteusin, RiPP regulation element (RRE), ribosomal synthesized and post-translationally modified clusters (RiPP), non-alpha poly-amino acid (NAPA), ribosomal peptide synthase clusters (RPS), siderophore, linear azoline containing peptides (LAP) and linaridin were detected only in few species (**Figure S6**). These metabolites could inhibit the growth of other organisms by blocking the biologically important processes (Oakley 2013). Our findings based on the genomic studies could be helpful in the search for secondary metabolites and their functions in this group of bacteria.

**Fig. 5.**
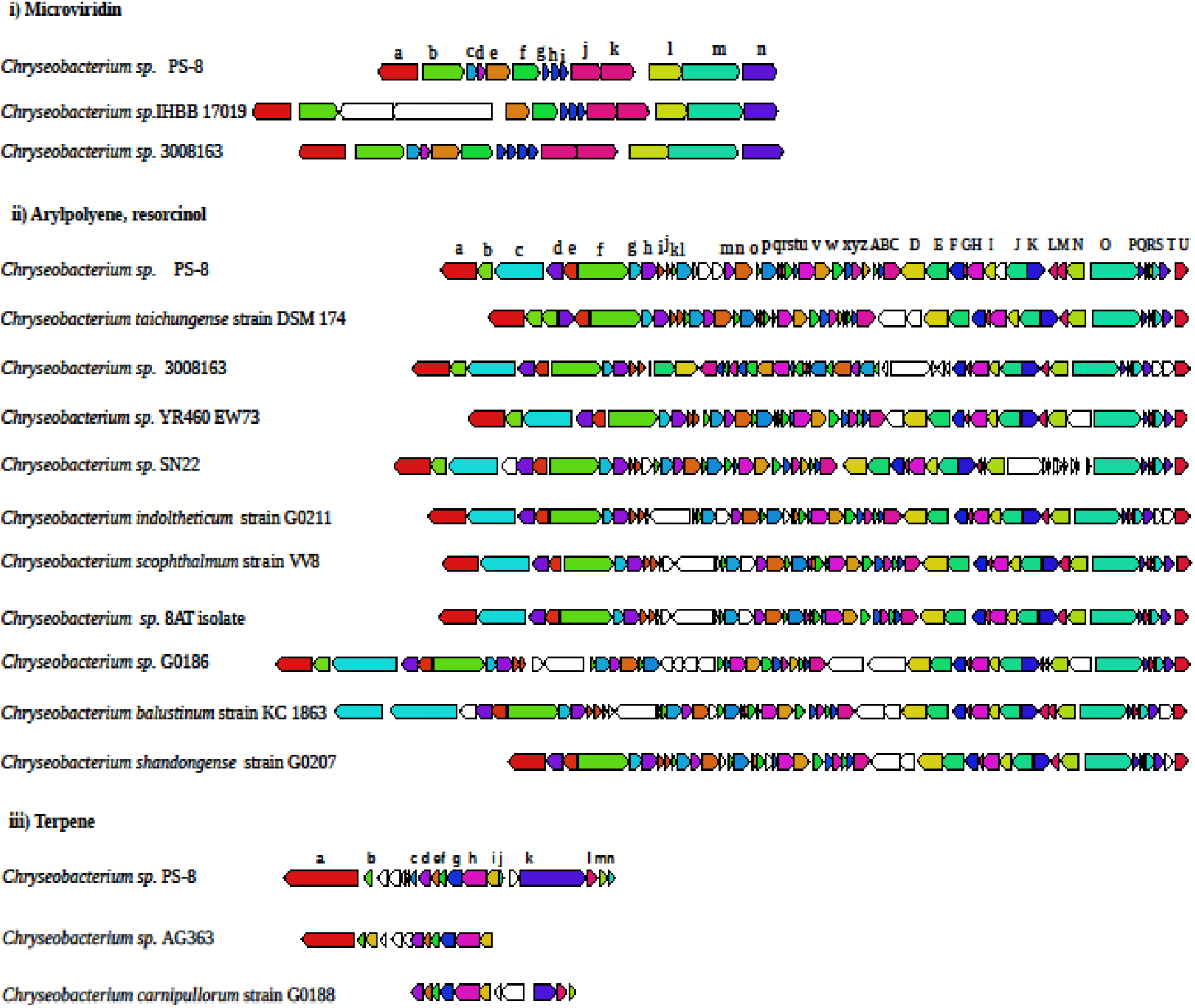
The secondary metabolite biosynthetic gene clusters (BGCs) predicted in the species of *Chryseobacterium*. The length of each horizontal bar corresponds with the number of BGCs. **i),** Microviridin related genes obtained in the *Chryseobacterium* sp PS-8 from this study and NCBI database. a, carboxypeptidase regulatory-like domain-containing protein; b, mechanosensitive ion channel; c, DUF2085 domain-containing protein; d, hypothetical protein; e, pyridoxine 5’-phosphate synthase; f, alpha/beta fold hydrolase; g, microviridin/ marinostatin family tricyclic proteinase inhibitor; h, microviridin/marinostatin family tricyclic proteinase inhibitor; I, microviridin/marinostatin family tricyclic proteinase inhibitor; j, ATP-grasp ribosomal peptide maturase; k, MvdD family ATP-grasp ribosomal peptide maturase; l, NAD-dependent epimerase/dehydratase family protein; m, long-chain fatty acid--CoA ligase; n, diphosphomevalonate decarboxylase. **ii),** Resorcinol and polyene related genes obtained in the *Chryseobacterium* sp PS-8 from this study and NCBI database. a, type I DNA topoisomerase; b, hypothetical protein; c, T9SS type A sorting domain-containing protein; d, EpsG family protein; e, hypothetical protein; f, MMPL family transporter; g, dialkylresorcinol condensing enzyme DarA; h, beta-ketoacyl-ACP synthase III; I, hypothetical protein; j, ABC transporter permease; k, hypothetical protein; l, BtrH N-terminal domain-containing protein; m, ABC transporter ATP-binding protein; n, ABC transporter permease; o, acyl-CoA thioesterase; p, beta-ketoacyl synthase; q, addiction module family protein; r, type II toxin-antitoxin system RelE/ParE family toxin; s, 3 oxoacyl-ACP synthase; t, acyl carrier protein; u, beta-ketoacyl-[acyl-carrier-protein] synthase family protein; v, beta-ketoacyl synthase chain length factor; w, polysaccharide deacetylase family protein; x, outer membrane lipoprotein carrier protein LolA; y, hypothetical protein; z, PorT family protein; A, 3-hydroxyacyl-ACP dehydratase; B, hypothetical protein; C, DUF2062 domain-containing protein; D, acyl-CoA--6-aminopenicillanic acid acyl-transferase; E, NAD(P)/FAD-dependent oxidoreductase; F, lipid A biosynthesis acyltransferase; G, acyl carrier protein; H, beta-ketoacyl-[acyl-carrier-protein] synthase family protein; I, 3-oxoacyl-ACP reductase FabG; J, aromatic amino acid lyase; K, hypothetical protein; L, hypothetical protein; M, hypothetical protein; N, AMP-binding protein; O, isoleucine--tRNA ligase; P, TraR/DksA family transcriptional regulator; Q, rhomboid family protein; R, hypothetical protein; S, lipoprotein signal peptidase; T, YdcF family protein; U, tryptophan--tRNA ligase. **iii),** Terpene related genes obtained in the genome of *Chryseobacterium* sp PS-8 from this study and NCBI database. a, CusA/CzcA family heavy metal efflux RND transporter; b, hypothetical protein; c, rhodanese-like domain-containing protein; d, lycopene cyclase domain-containing protein; e, sterol desaturase family protein; f, SRPBCC family protein; g, phytoene/squalene synthase family protein; h, phytoene desaturase; I, winged helix DNA-binding protein; j, hypothetical protein; k, DUF11 domain-containing protein; l, hypothetical protein; m, hypothetical protein; n, helix-turn-helix domain-containing protein.

### Description of *Chryseobacterium indicum* sp. nov

*Chryseobacterium indicum* sp. nov. (in’di.cum. L. neut. adj. *indicum*, of or belonging to India, the country where the type strain was isolated).

Strain PS-8 is Gram-negative, aerobic, rod-shaped approximately 0.5–0.75 μm wide and 0.8 −2.0 μm long. Colonies of strain PS-8 appeared as yellowish-orange colour. Growth occurred at 10-40 °C and pH 5–10. Optimal growth was observed at 28–30 °C and pH 6.5-7. It can grow at 1% (w/v) of NaCl. Positive for catalase, oxidase, hydrolysis of gelatin, DNA, casein, aesculin, tween 40 and tween 80 and negative for arginine dihydrolase, lysine decarboxylase, production of H_2_S, the activity of galactopyranosidase, ornithine decarboxylase, tryptophan deaminase, Voges-Proskauer, urease, citrate utilization, hydrolysis of starch, and cellulose, production of acetoin, nitrate, and nitrite reduction. Acid is produced from sucrose, arabinose and glucose. Predominant fatty acids are 15:0 iso, 17:0 iso 3OH, 15:0 iso 3OH and 11:0 anteiso. Polar lipids included phosphatidylethanolamine (PE) and amino lipids (AL). The G+C content is 36.4%.

The type strain is PS-8^T^ (= TBRC 15233^T^ = NBRC 115235 ^T^), which was isolated from the skin of freshwater pufferfish.

## MATERIALS AND METHODS

### Bacterial strain and growth medium

*Chryseobacterium* strain PS-8 was isolated from the skin of freshwater pufferfish as described earlier (Das *et al.* 2021). A colony with yellowish-orange colour designated strain PS-8 was selected for further analyses.

### Phenotypic characterization

Cell morphology was examined using a confocal microscope (TCS SP5, Leica). Motility was tested by the hanging drop method (Bernardet *et al.* 2002). Gram staining was done using a commercial kit (Becton–Dickinson). Oxidase activity was assayed commercial kit (Hi-Media). Hydrogen peroxide (10%, v/v) was used for the catalase activity. Anaerobic growth was tested using the BD GasPak EZ system (Becton Dickinson). Growth at different pH ranges (4.0–13.0; in steps of 0.5 pH units) and temperatures (4–50 °C; in steps of 2 °C) was determined in nutrient broth (NA, Difco). Growth at different temperatures was monitored by measuring OD_600_ with a spectrophotometer (CARY 300, UV–Vis spectrophotometer, Varian). The growth in different concentrations of NaCl (0–15 % (w/v) was determined in MM63 minimal media and incubated at 28 °C for 2 days (Jiang *et al.* 2014). Phenotypic characteristics such as Voges Proskauer test, indole, methyl red, H_2_S production, citrate utilization, nitrate reduction, phenylalanine deaminase, hydrolysis of esculin, lysine decarboxylase, ornithine decarboxylase, arginine dihydrolase, urease, casein, DNA, gelatin, starch and Tween 20 and 80 were tested according to Mata *et al.* (2002). Phenotypic characterization was also performed using API 20 NE, API 20E, API ZYM (bioMérieux) and Biolog GNII microplates. The haemolytic activity was tested on Columbia blood agar (Patra *et al.* 2007).

### Chemotaxonomic analysis

For lipid analysis, cells were grown in nutrient broth (Difco) to mid-exponential phase on a rotary shaker at 28 oC. 200 mg of lyophilized cells were used for polar lipid extraction as described by Bligh and Dyer (1959) and separated by two-dimensional TLC (silica gel 60 F254, catalogue number 1.05554.0007; Merck). The solvent system used has been described earlier (Kumari *et al.* 2014). Diaminopimelic acid (DAP) and cell wall sugar were detected following the method of Staneck and Roberts (1974). For fatty acids, cells grown on nutrient agar (Difco) plates at 28 °C for three days, saponified, and analysed using the Sherlock Microbial identification system (MIDI, Microbial ID), consisting of a model 6890 N gas chromatograph (Agilent).

### Whole genome sequencing and annotation

The genomic DNA was isolated using the methods of Sambrook and Russel (2001). DNA concentration was measured using NanoDrop 8000 Spectrophotometer (Thermo Scientific). Both Illumina and Oxford Nanopore sequencing platforms were used to generate the genome sequence. Illumina short-read DNA sequencing was done as described earlier (Wick *et al.* 2017). For long-read, a genomic library was prepared using the Nanopore Ligation Sequencing Kit (SQK-LSK109; Oxford Nanopore, Oxford, UK). Assembly and annotation were done as described earlier (Das *et al.* 2022). The gene ontology (G.O.) study was performed using the Blastp program (https://www.biobam.com/omicsbox) implemented in Blast2GO tool (Götz *et al.* 2008) for cataloguing gene function.

### Comparative genomics and phylogenetic analysis

*In silico* DDH similarity and AAI, ANI was estimated following the methods of (Meier-Kolthoff *et al.* 2013). For phylogenomic analysis, 73 whole-genome sequences of *Chryseobacterium* type species with more than 95% genome completeness were retrieved from the NCBI database. The core genes were extracted by the UBCG pipeline (Na *et al.* 2018). The maximum-likelihood tree was reconstructed with the GTR model (Stamatakis 2014). Further, the non-recombinant core-genome based phylogenetic tree was constructed by Mateo-Estrada *et al.* (2019).

### Prediction of virulence genes and secondary metabolite

Blast and reciprocal blast (blastp) (Altschul *et al.* 1990) were performed (query coverage, percentage identity >=40%) with clinically verified core data-set (3580 genes) of virulent factor database (VFDB) (Chen *et al.* 2005) to retrieve the putative virulence genes in PS-8 and the type strains compared in this study. Cladogram-heatmap was generated using the ETE-3 package of the python program. Hits with either coverage or identity lower than 40% were not considered. The secondary metabolites produced by these bacterial species was predicted using antiSMASH 5.0 (Blin *et al.* 2019) and BiG-SCAPE (Navarro-Muñoz 2020) software tools with default parameters. The genome mining algorithms employed for the secondary metabolite prediction are derived from small-molecule-producing bacteria, like *Myxobacteria* and *Actinobacteria* (Scherlach and Hertweck 2021; Baral *et al.* 2018).

### Data availability

The GenBank/EMBL/DDBJ accession number for the genome and 16S rRNA gene sequences of *Chryseobacterium indicum* strain PS-8 are <u>JACSGT000000000</u> and <u>MZ305332</u>, respectively.

## ACKNOWLEDGEMENTS

The author M.P.V. and L.D. acknowledges the Department of Biotechnology, Government of India and Council of Scientific and Industrial Research (CSIR), New Delhi, Government of India for providing the research fellowship. This work was supported by the funding received from the Department of Biotechnology, Government of India (D.O.No. BT/BI/04/058/2002 VOL-II) to SKD under the Distributed Information Sub-Center (DISC) project.

## Ethical approval

This study was carried out with approval from the Institutional Animal Ethics Committee (Letter No. V-311-MISC/2017-18/ILS/884).

## Authors contribution

S.K.D. developed the concept. M.P.V. and L.D. participated in laboratory experiments. M.P.V, L.D. and S.K.D. analysed the data and wrote the manuscript. All of us read and approved the final manuscript.

## Conflict of interest

The authors declare no conflicts of interest.

## Supplementary materials

**Supplementary Table S1.**
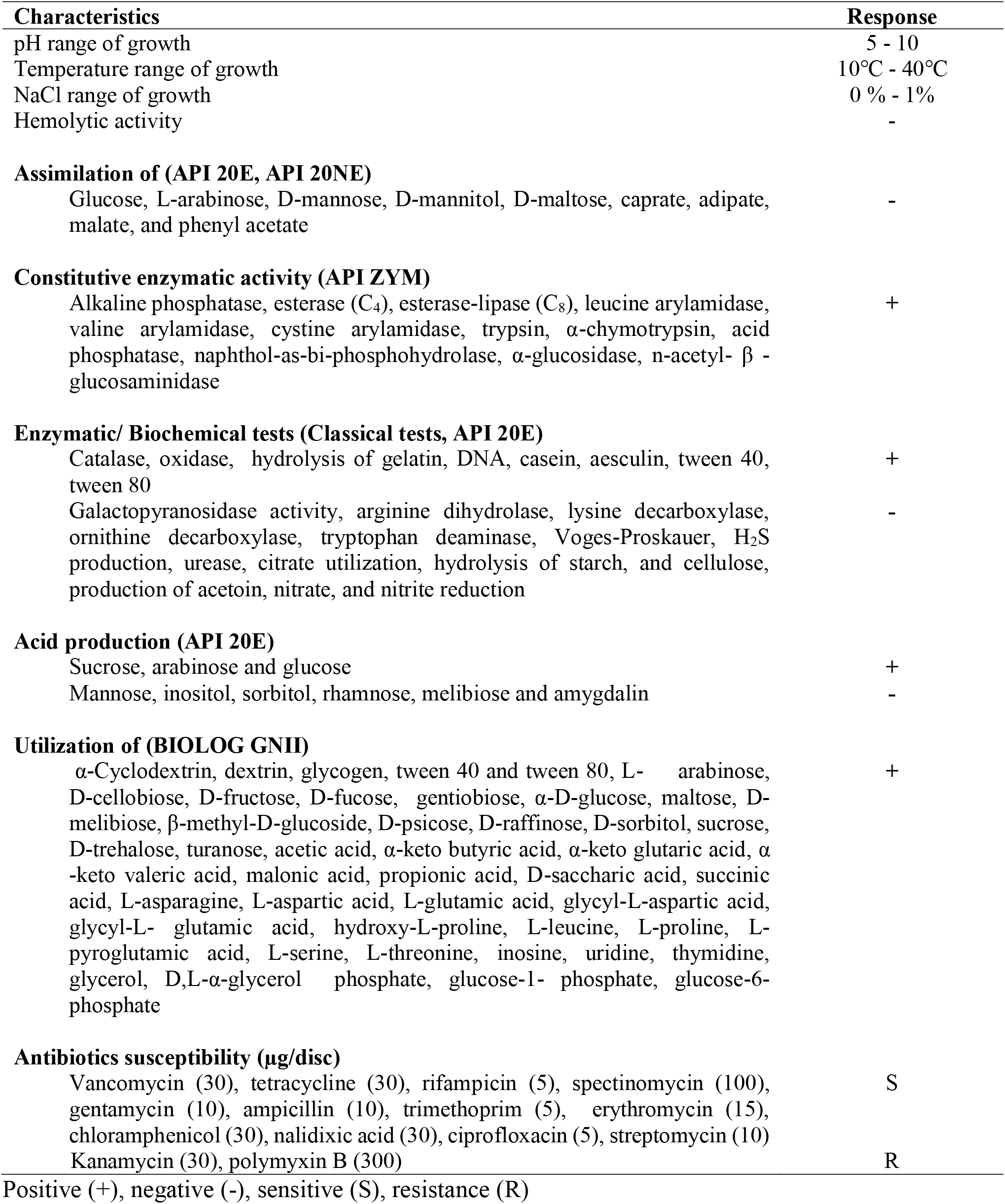
Phenotypic features of *Chryseobacterium* strain PS-8.

| Characteristics | Response |
| --- | --- |
| pH range of growth | 5 - 10 |
| Temperature range of growth | 10°C - 40°C |
| NaCl range of growth | 0 % - 1% |
| Hemolytic activity | - |
| **Assimilation of (API 20E, API 20NE)** |  |
| Glucose, L-arabinose, D-mannose, D-mannitol, D-maltose, caprate, adipate, malate, and phenyl acetate | - |
| **Constitutive enzymatic activity (API ZYM)** |  |
| Alkaline phosphatase, esterase (C<sub>4</sub>), esterase-lipase (C<sub>8</sub>), leucine arylamidase, valine arylamidase, cystine arylamidase, trypsin, α-chymotrypsin, acid phosphatase, naphthol-as-bi-phosphohydrolase, α-glucosidase, n-acetyl- β -glucosaminidase | + |
| **Enzymatic/ Biochemical tests (Classical tests, API 20E)** |  |
| Catalase, oxidase, hydrolysis of gelatin, DNA, casein, aesculin, tween 40, tween 80 | + |
| Galactopyranosidase activity, arginine dihydrolase, lysine decarboxylase, ornithine decarboxylase, tryptophan deaminase, Voges-Proskauer, H<sub>2</sub>S production, urease, citrate utilization, hydrolysis of starch, and cellulose, production of acetoin, nitrate, and nitrite reduction | - |
| **Acid production (API 20E)** |  |
| Sucrose, arabinose and glucose | + |
| Mannose, inositol, sorbitol, rhamnose, melibiose and amygdalin | - |
| **Utilization of (BIOLOG GNII)** |  |
| α-Cyclodextrin, dextrin, glycogen, tween 40 and tween 80, L- arabinose, D-cellobiose, D-fructose, D-fucose, gentiobiose, α-D-glucose, maltose, D-melibiose, β-methyl-D-glucoside, D-psicose, D-raffinose, D-sorbitol, sucrose, D-trehalose, turanose, acetic acid, α-keto butyric acid, α-keto glutaric acid, α -keto valeric acid, malonic acid, propionic acid, D-saccharic acid, succinic acid, L-asparagine, L-aspartic acid, L-glutamic acid, glycyl-L-aspartic acid, glycyl-L- glutamic acid, hydroxy-L-proline, L-leucine, L-proline, L-pyroglutamic acid, L-serine, L-threonine, inosine, uridine, thymidine, glycerol, D,L-α-glycerol phosphate, glucose-1- phosphate, glucose-6-phosphate | + |
| **Antibiotics susceptibility (μg/disc)** |  |
| Vancomycin (30), tetracycline (30), rifampicin (5), spectinomycin (100), gentamycin (10), ampicillin (10), trimethoprim (5), erythromycin (15), chloramphenicol (30), nalidixic acid (30), ciprofloxacin (5), streptomycin (10) | S |
| Kanamycin (30), polymyxin B (300) | R |

Positive (+), negative (-), sensitive (S), resistance (R)

**Supplementary Table S2.**
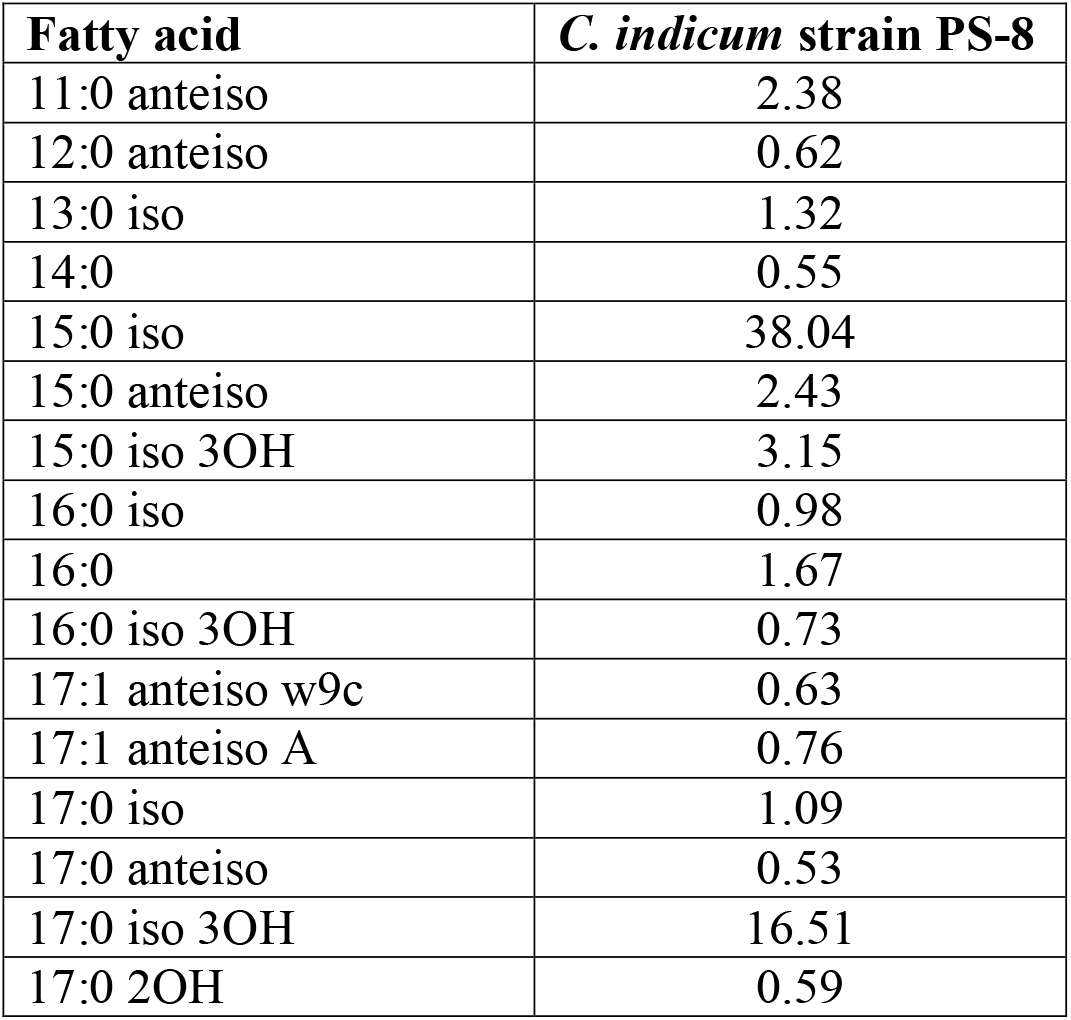
Fatty acid profile of *Chryseobacterium* strain PS-8.

## Supplementary figures

**Figure S1.**
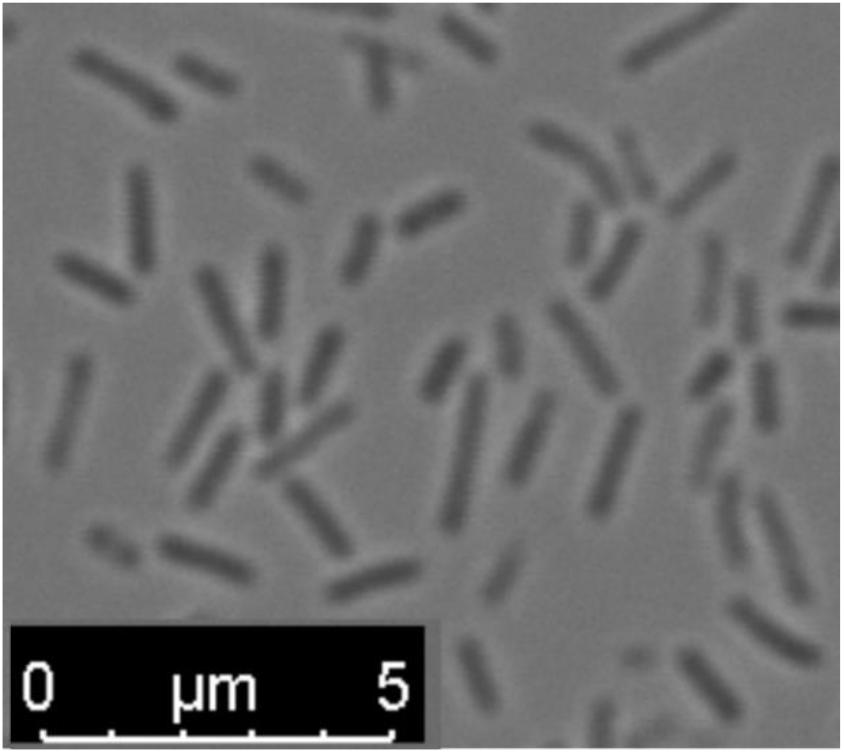
Morphology of *Chryseobacterium* strain PS-8.

**Figure S2.**
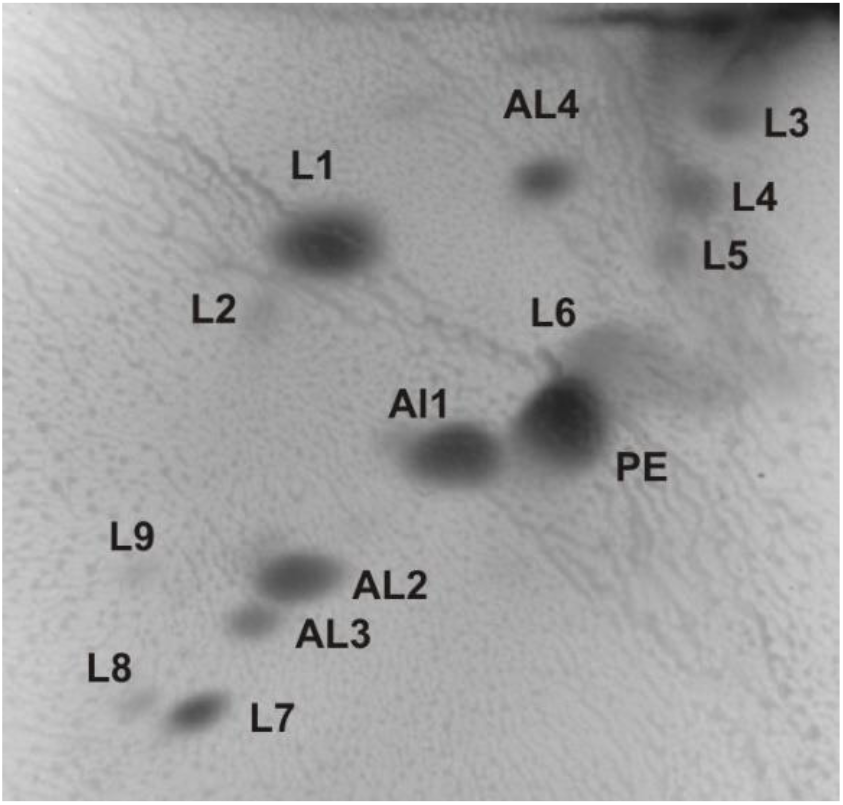
Two-dimensional thin-layer chromatographic profile of polar lipids of *Chryseobacterium* strain PS-8. Abbreviations: Phosphatidylethanolamine (PE), amino lipid (AL) and unknown lipid (L1-L9).

**Figure S3.**
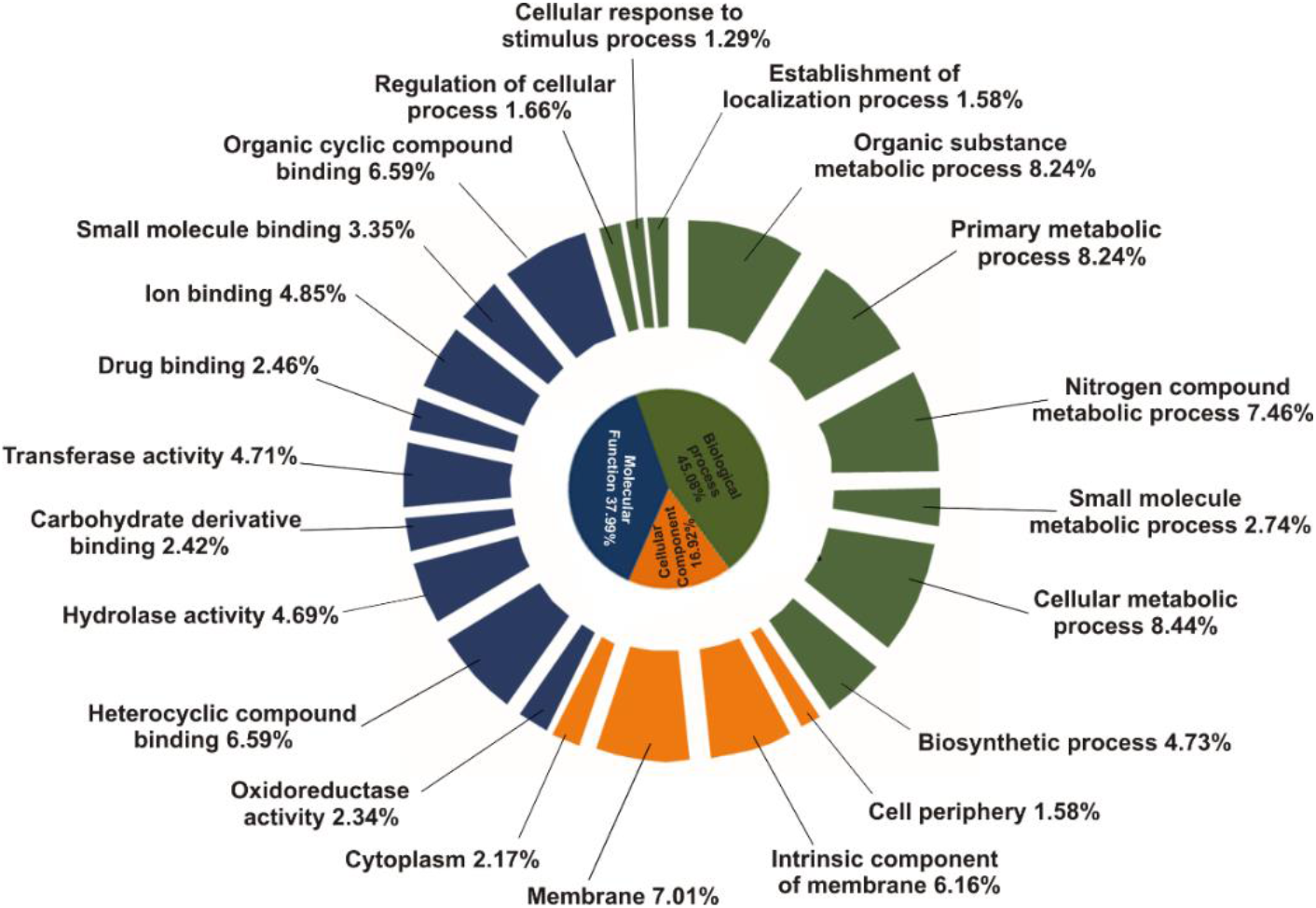
Gene ontology (GO) association of predicted protein-coding genes of *Chryseobacterium* strain PS-8.

**Figure S4.**
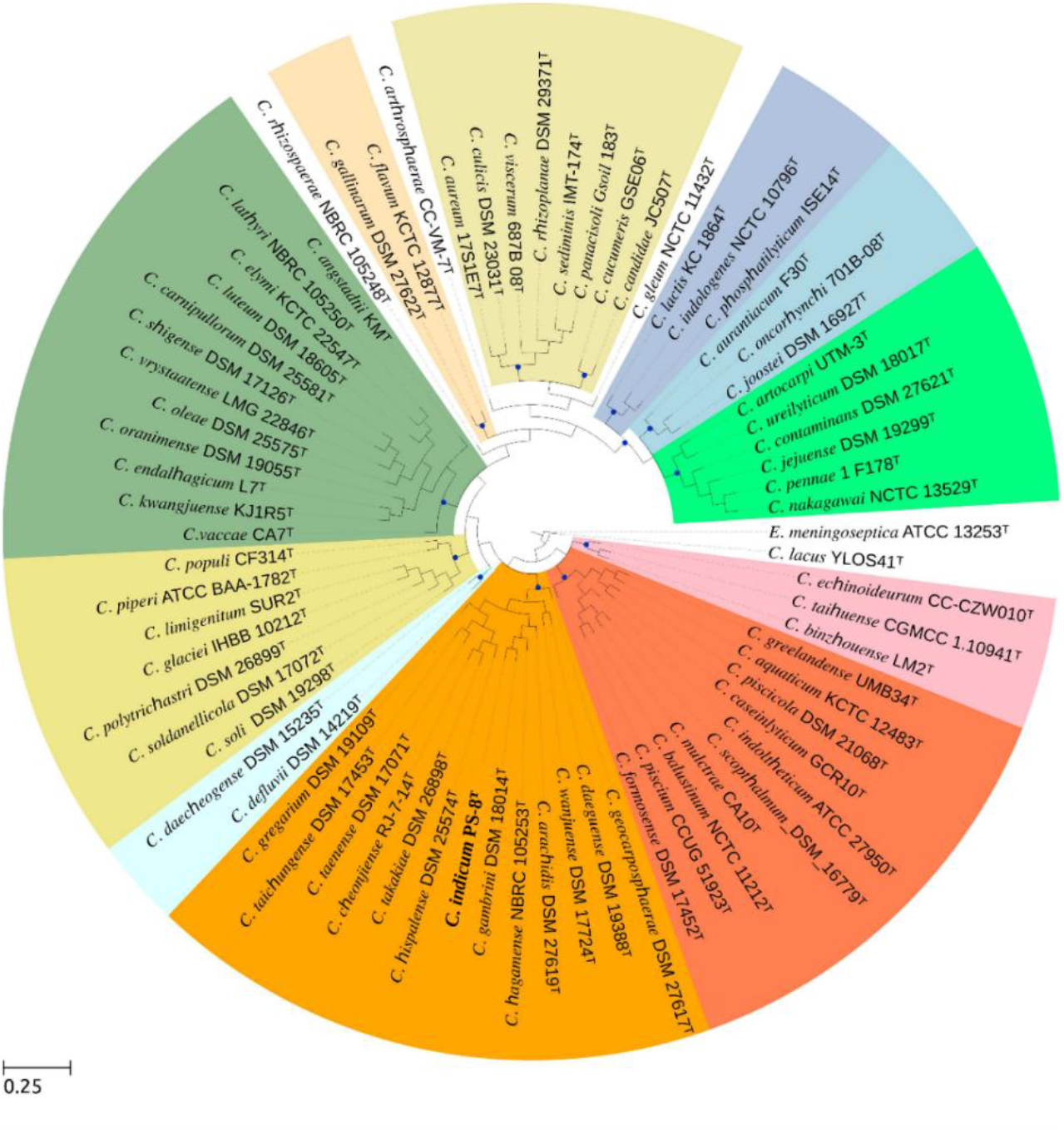
Maximum-likelihood phylogenetic tree based on the alignment of non-recombinant genes of strain PS-8 and 73 type species of *Chryseobacterium*. The tree was constructed with ETE-3 python package and rooted using *Elizabethkingia meningoseptica* ATCC 13253T as the out group. Bar, 0.25 substitutions per nucleotide position.

**Figure S5.**
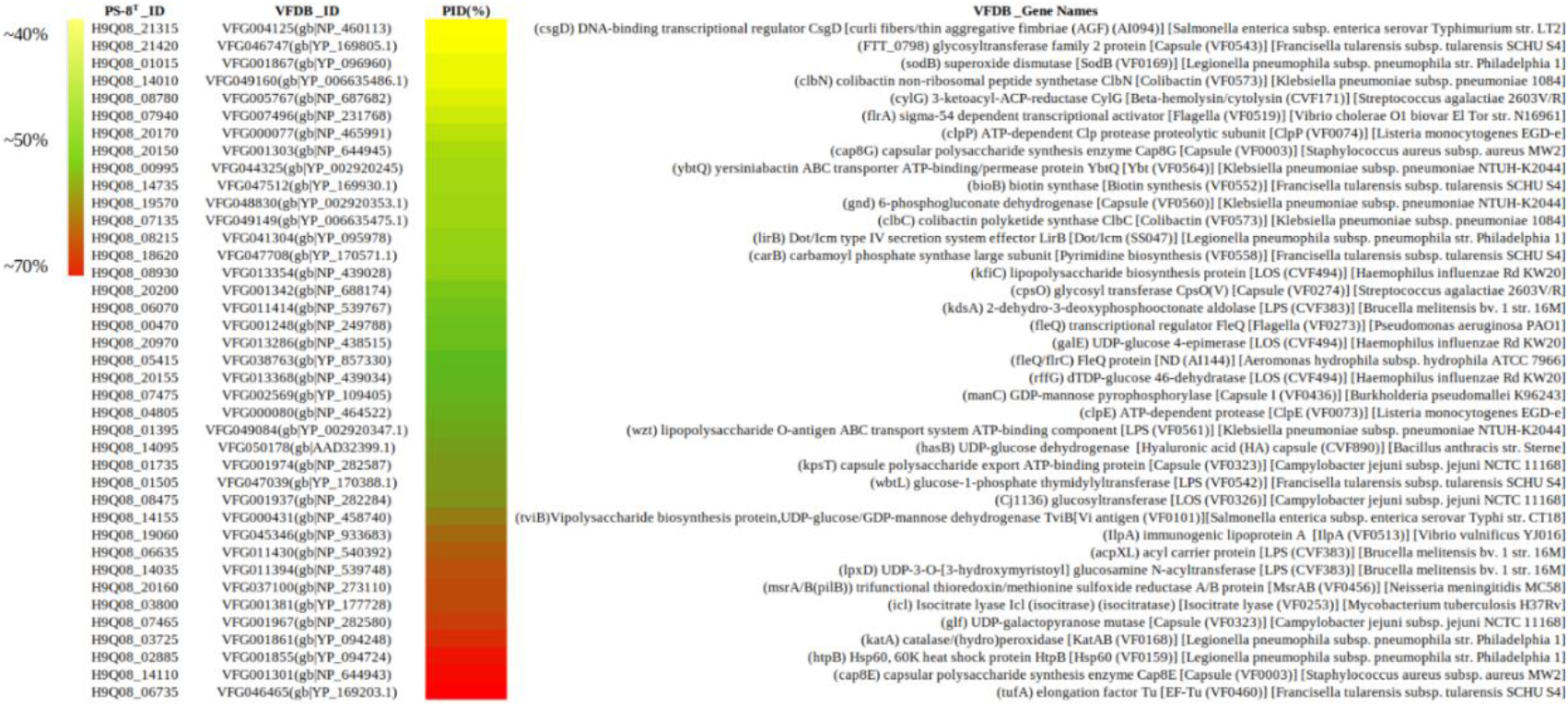
Virulence associated proteins identified in strain PS-8. Gene id was derived from the NCBI gene bank file and VFDB id from the virulence factor database.

**Figure S6.**
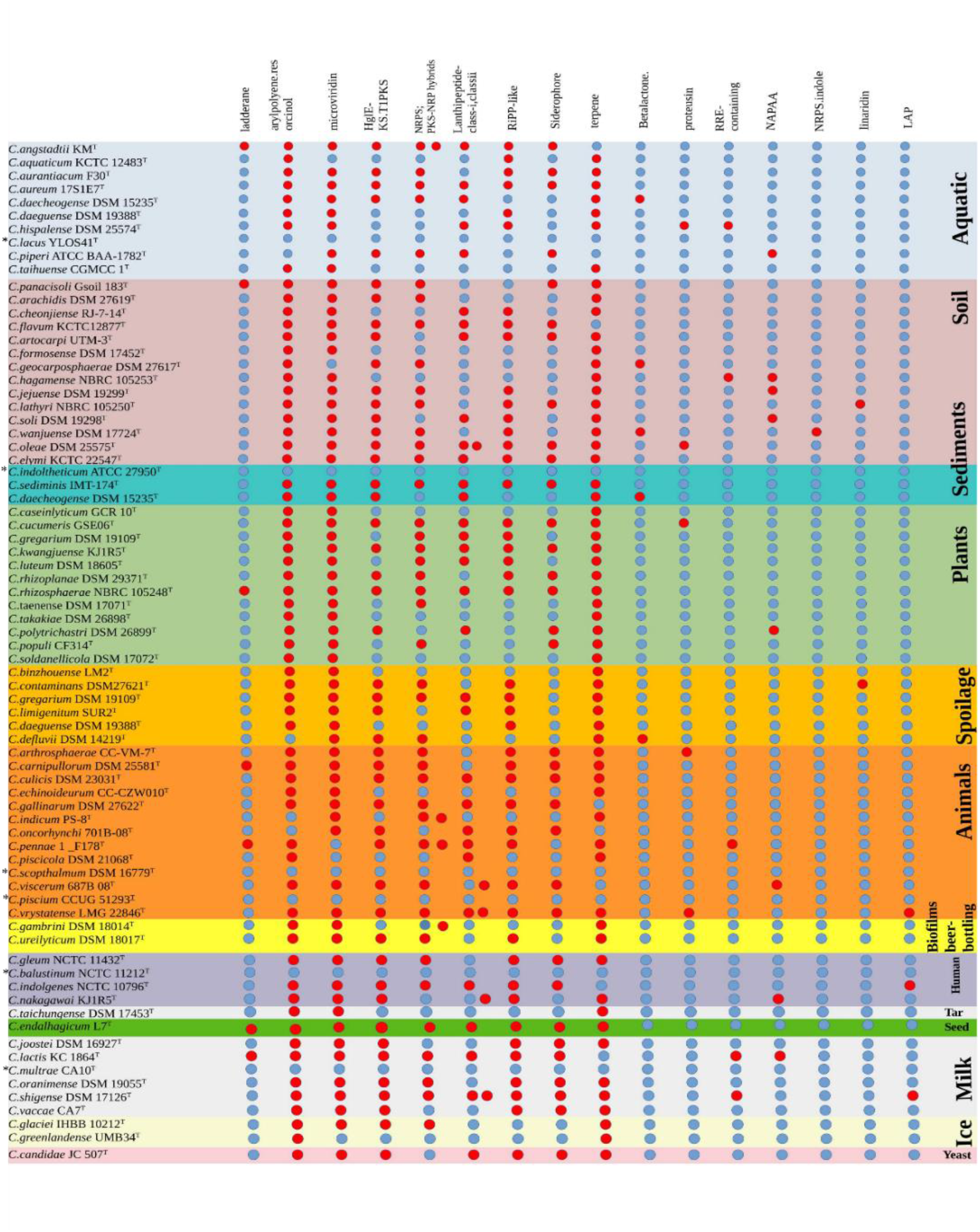
Prediction of secondary metabolites production for each *Chryseobacterium* type strain. The red colour indicates presence, and the grey colour indicates absence. *Secondary metabolites not detected.

## Notes

### Competing Interest Statement

The authors have declared no competing interest.

